# The Feather Epithelium Contributes to the Dissemination and Ecology of clade 2.3.4.4b H5 High Pathogenicity Avian Influenza Virus in Ducks

**DOI:** 10.1101/2023.07.26.550633

**Authors:** Nicolas Gaide, Fabien Filaire, Kateri Bertran, Manuela Crispo, Malorie Dirat, Aurélie Secula, Charlotte Foret-Lucas, Bruno Payré, Albert Perlas, Guillermo Cantero, Natàlia Majó, Sébastien Soubies, Jean-Luc Guérin

## Abstract

Immature feathers are known replication sites for high pathogenicity avian influenza viruses (HPAIVs) in poultry. However, it is unclear whether feathers play an active role in viral transmission. This study aims to investigate the contribution of the feather epithelium to the dissemination of clade 2.3.4.4b goose/Guangdong/1996 lineage H5 HPAIVs in the environment, based on natural and experimental infections of domestic ducks. During the 2016-22 outbreaks, H5 HPAIVs exhibited persistent and marked feather epitheliotropism in naturally infected commercial ducks. Infection of feathers resulted in epithelial necrosis, disruption, and the production and release of infectious virions. Viral and feather antigens colocalized in dust samples obtained from poultry barns housing naturally infected birds. In summary, the feather epithelium contributes to viral replication, and it is a likely source of environmental infectious material. This underestimated excretion route could greatly impact the ecology of HPAIVs, facilitating airborne and preening-related infections within a flock, and promoting prolonged viral infectivity and long-distance viral transmission between poultry farms.

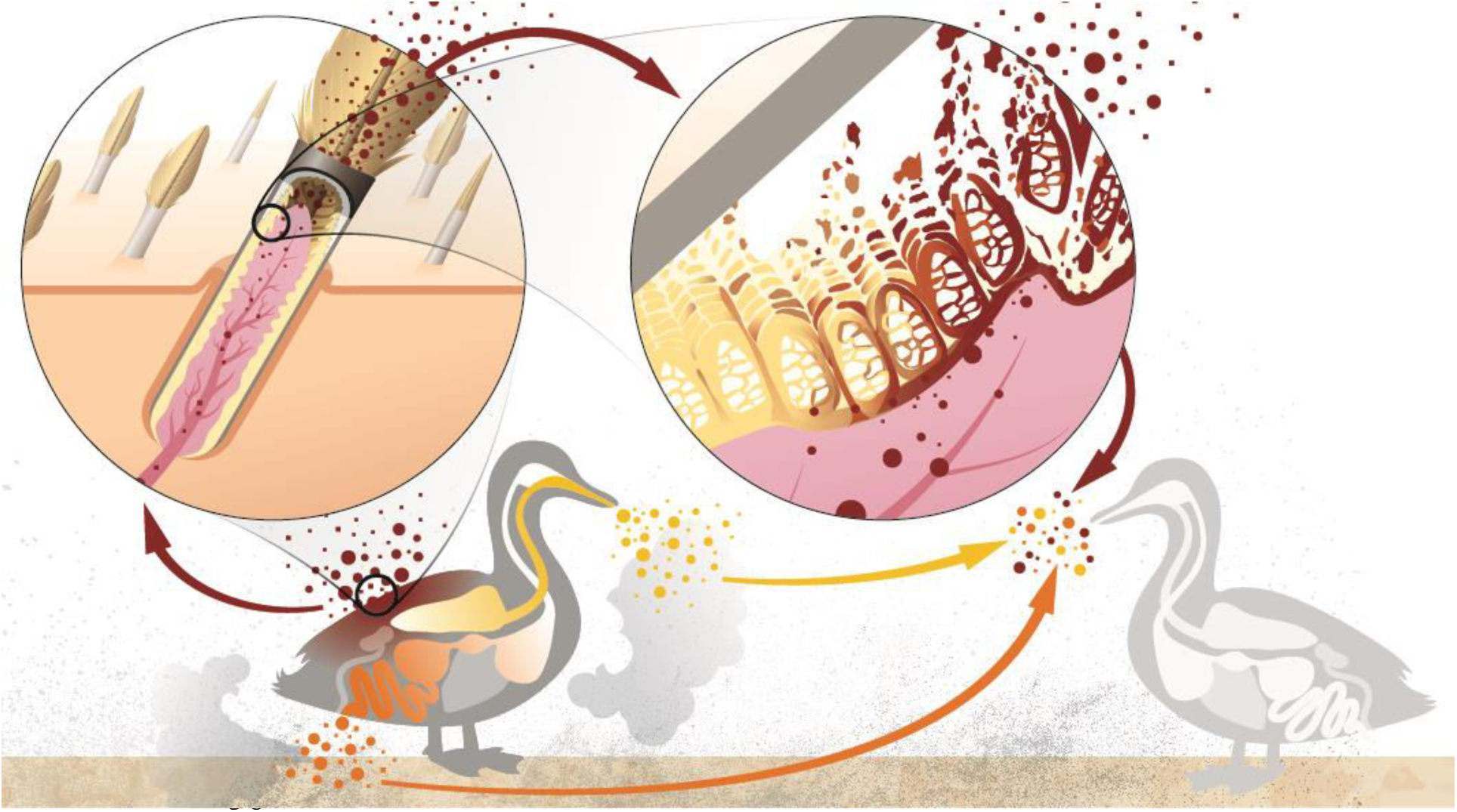

## Introduction

High pathogenicity avian influenza viruses (HPAIVs) of clade 2.3.4.4b goose/Guangdong/1996 (Gs/GD) H5 lineage have greatly impacted both the poultry industry and wild bird populations during the last few years, resulting in large-scale culling of infected flocks and mass mortality of endangered species worldwide^1, 2^. Ducks, in particular, play a crucial role in the transmission of HPAIVs due to the shedding of large amounts of infectious material during the pre-symptomatic phase, which can last several days. The main shedding routes include digestive and respiratory tracts^3–5^. Therefore, to develop effective control measures, it is essential to properly understand the mechanisms underlying HPAIVs infection and dissemination in ducks. In commercial ducks, infection with clade 2.3.4.4b H5 HPAIVs leads to neurological disease and mortality. Typical pathological findings include acute encephalitis, myocarditis, and pancreatitis. However, since these viruses replicate systemically, they can be detected in many other tissues^3, 6^.

A key feature of viral infection is tropism, which refers to the ability of a virus to infect and replicate in specific cells or tissues. Viral tropism plays a critical role in viral ecology since the range of infected tissues and organs determines clinical signs, mean death time and viral excretion routes^6^. Growing feather follicles have been identified as a site of H5 and H7 HPAIVs replication in ducks^7–9^ and other gallinaceous species^10–12^. Feathers are abundant external cutaneous appendages implicated in a variety of functions, including flight, insulation, and communication^13^. Immature feathers are growing tubular structures budding from the skin, covered by a protective external sheath and nourished by a vascularized dermal pulp (**supplementary data 1**). The feather epithelium grows and differentiates into barbs and barbules. Upon maturation, the external sheath opens and the epithelium comes into contact with the air^14^. Experimental data show that infected feathers pulp are a site of productive viral replication, and feathers obtained from infected duck carcasses contain infectious material for up to 160 days and 15 days post-mortem at 4°C and 20°C, respectively^12, 15^. Furthermore, the stability of avian influenza viruses has been reported to increase with preening oil, which is secreted by the uropygial gland and used for preening behavior^16^.

Dust found in poultry houses is composed of diverse particles in terms of size, density, and shape, originating from feed, feces, skin, feather and litter debris, microorganisms, and insect parts^17–19^. Feathers are reported to account for 10% of dust found in commercial layer chicken houses^17, 18^. Not only is dust known to contribute to viral dispersal within farms, but also inhalable particles (<100 µm) can lead to bird-to-bird airborne transmission^19–23^. While dust is a recognized source of pathogens^19^, the specific contribution of feathers in infectious viral shedding, as well as viral environmental spread and ecology, remains unclear.

This study aims to investigate the role of feather epitheliotropism in the environmental dissemination of clade 2.3.4.4b H5 HPAIVs, combining natural and experimental infections in commercial ducks.

## Material and Methods

### Field cases

Eight flocks of commercial ducks (two flocks of Muscovy ducks [*Cairina moschata*] and six flocks of mule ducks [*Cairina moschata x Anas platyhrynchos*]) positive for H5 HPAIVs by official testing during the 2016-22 HPAIVs outbreaks in France were included in the study. Pathological investigations were conducted on a total of 36 dead animals, naturally infected with H5N8 2016-17 (*n=8*), H5N8 2020-21 (*n=10*), and H5N1 2021-22 (*n=18*) HPAIVs. All samples were obtained under the supervision of the French Official Veterinary Services and in compliance with the French legislation on veterinary practices and notifiable diseases. At necropsy, feathered skin (including wing feathers, caudal tract, and capital tract) was collected and fixed in 10% neutral buffered formalin for histopathological examination and viral *in situ* detection. Virological titration was conducted on two flocks of ducks naturally infected with H5N1 2021-22 HPAIV (5 ducks per flock). Additionally, feathered skin from dead chicken (4 flocks, *n=3-5*), turkey (1 flock, *n=4*), quail (1 flock, *n=5*), geese (1 flock, *n=6*) and wild swans (3 birds) naturally infected with H5 HPAIV from 2016 to 2022 was collected, treated, analyzed similarly to duck and provided supplementary data.

### Experimental infection

Fifty 5-week-old domestic male mule ducks were obtained from a commercial producer (GALLSA, Tarragona, Spain). Ducks were challenged by intrachoanal inoculation with 5 log_10_ 50% egg infectious doses (EID_50_) of either A/mulard duck/France/171201g/2017(H5N8) (H5N8/2017) reverse genetics-engineered virus (accession numbers MK859904 to MK859911)^24^ or A/Mule duck/France/20320/2020(H5N8) (H5N8/2020) (accession numbers MZ166297 to MZ166304), and monitored for 14 days post-inoculation (dpi). Groups included sham birds (n=3), inoculated birds (n=16) and contact birds placed 24 hours post-inoculation (n=6), for each viral strain. Feather pulp samples were collected on 10 inoculated and 6 contact birds at 0, 2, 4, 7, 10 and 14 dpi for viral RNA detection. Complete necropsies were performed at 3 and 5 dpi for each virus on 3 inoculated birds and on 3 sham birds. At necropsy, sections of feathered skin, including capital and caudal tracts^9^, as well as detached immature wing feathers were collected for histopathology and stored in 10% neutral buffered formalin. All procedures involving birds were reviewed and approved by the IRTA (#258-2021) and the Catalan Government (#11467) Ethics and Animal Experimentation Committees, subject to national and European regulations. All procedures involving virus were performed in biosafety level-3 (BSL-3) laboratory and animal facilities at IRTA-CReSA, in accordance with procedures approved by IRTA Biosafety Committee (#59-2021).

### Viral molecular detection

Total RNA was extracted from experimental and selected field samples using the magnetic bead-based ID Gene Mag Fast Extraction Kit and IDEAL 32 extraction robot (Innovative Diagnostics, Grabel, France) according to the manufacturer’s instructions. A one-step, real-time reverse transcription quantitative PCR (rRT-qPCR) was then performed using the Influenza H5/H7 Triplex kit (Innovative Diagnostics, Grabel, France).

### Histology and viral *in situ* detection

Fixed tissue samples were paraffin-embedded and cut at 3 µm. Sections were stained with hematoxylin and eosin (H&E) for histopathological analysis and anti-influenza A virus nucleoprotein immunohistochemistry (anti-IAV NP IHC) using anti-NP influenza A HB65 antibodies (NP, HB65, lot 758121J1, Biozol, Eching, Germany). Briefly, the IHC protocol included an antigen retrieval step with 0.05% pronase applied for 10 min at 37°C, a peroxidase blocking step of 5 min at room temperature (S2023; Agilent) followed by saturation of nonspecific binding sites with normal goat serum (X0907; Agilent) applied for 25 min at room temperature, and overnight incubation with anti-IAV NP antibody (1:2000 dilution) at 4°C. Signal amplification and revelation were assessed using the EnVision FLEX system and 3,3’-diaminobenzidine (DAB) revelation according to the manufacturer’s recommendations. Avian Influenza *Matrix* gene (*M* gene) RNAscope *in situ* hybridization (RNAscope ISH) was performed on selected samples, as previously described^25^. Viral antigen detection was determined for epithelial and mesenchymal tissues of growing feathers, resting feather epidermis and dermis^26^. Results were expressed as frequency (number of positive birds / number of total birds *100) and distribution was determined according to Landman (2021) for each flock^27^.

### Transmission Electron Microscopy

Ultrastructural analyses were performed on samples obtained from an experimentally infected duck with H5N8/2017, which was selected based on histopathological lesions and viral antigen detection. Growing feathers from the caudal tract stored in formalin were cut (6 transversal sections), immersed in 2% glutaraldehyde in Sorensen’s phosphate buffer (0.1 mol/L, pH = 7.4) for 1 hour, and washed with Sorensen’s phosphate buffer for 12 hours. Then samples were incubated with 1% OsO_4_ in Sorensen’s phosphate buffer (0.05 mol/L, glucose 0.25 mol/L, OsO_4_ 1%) for 1 hour, dehydrated in an ascending ethanol series until ethanol 100° and then with propylene oxide, and embedded with epoxy resin (EMBed 812). After 48h of polymerization at 60°C, ultrathin sections (70 nm) were mounted on 100 mesh collodion-coated copper grids and post-stained with 3% uranyl acetate in 50% ethanol and with 8.5% lead citrate before being examined on a HT 7700 Hitachi electron microscope at an accelerating voltage 80 KV.

### Dust morphological and immunofluorescent study

Dust samples were collected from four H5N1 2021-22 HPAIV-positive farms using a dry cyclonic air sampler Coriolis Compact (Bertin Technologies, https://www.bertin-instruments.com) for 20 min at 50 L/min. Samples were resuspended in 1 mL of phosphate buffered saline (PBS). Dust microscopic examination was conducted following an agar-based method, with a similar approach to the cell block technique used in cytopathology^28^. Briefly, 100 µL were centrifuged for 10 min at 300 g. The supernatant was removed and the pellet, consisting of dust, was resuspended in 50 µL of PBS and then 100 µL of Histogel® heated at 60 +/- 5°C (Thermo Scientific Richard-Allen Scientific, Waltham, MA). Dust blocks were formalin-fixed, paraffin-embedded and serially cut at 3 µm. One section was stained with H&E for histological analysis. On a second section, a double immunofluorescence (IF) was performed using non-commercial polyclonal rabbit primary antibodies targeting corneous beta-proteins (CBP) and mouse monoclonal anti-NP influenza A antibodies^29^.The antigenic retrieval and saturation steps were similar to those used in the anti-IAV NP IHC. Both antibodies were co-incubated at dilutions of 1/2000 (IAV NP) and 1/100 (CBP) overnight at 4°C. Anti-mouse 594 (Invitrogen, A11032) and anti-rabbit 488 (Jackson ImmunoResearch, 711-546-152) secondary antibodies were then used at a dilution of 1/500, incubated 1h at room temperature. Slides were mounted with Fluoromount G with DAPI (Thermo-Fischer Scientific, 00-4959-52). Image were collected using a Zeiss LSM 710 confocal microscope.

### Viral titration

Growing feathers collected from ten ducks belonging to two flocks (five ducks per flock) that tested positive for H5N1 during the 2021 outbreaks in France were used to determine the infectivity of the feather subcompartments through viral titration. Feather tips were sealed with heated paraffin, and the feather outer sheath was decontaminated by immersion in 70% ethanol for 5 min at room temperature, followed by a washing step in PBS with 0.5% bovine serum albumin (BSA). Feather outer sheath was then longitudinally opened to dissect both the distal epithelial fraction (whitish, matt, filamentous material) and the basal pulp (pink-red, soft, translucent and moist material). Epithelial samples were incubated in PBS-0.5% BSA for 30 min at room temperature. Subsequently, the PBS-0.5% BSA solution was replaced and vortexed for 30 seconds. Feather outer sheath (before and after decontamination), basal pulp, and distal epithelial fractions (with and without mechanical disruption) were subjected to viral titration. Titrations were performed using Madin-Darby canine kidney (MDCK) cells, followed by an immunoperoxidase monolayer assay. Results were expressed as log_10_ focus forming units per mL (FFU/mL) according to Matrosovitch *et al.*, 2006^30^.

### Statistics

Statistical analyses were performed using R Studio software (https://www.R-project.org) to compare viral titers among tissue compartments (feather outer sheath before and after decontamination, distal feather epithelium before and after vortexing, and basal pulp). Data for each sample were analyzed with a linear model that used the bird as a random parameter, and the tissue compartment and flock as fixed parameters.

## Results

### H5 HPAIVs lead to severe damage of growing feather epithelium

Growing feathers of domestic ducks naturally infected with H5 HPAIVs between 2016 and 2022 exhibited segmental and multifocal to diffuse epithelial necrosis (**Figure 1A**). Lesions were identified in several layers exhibiting different stages of differentiation, including the epidermal collar and early and late barb ridges, and were associated with epithelial disruption and loss of the original architecture (**supplementary data 2**). Epithelial necrosis and signs of dermal pulpitis, such as hyperemia, exudation, leukocytic infiltration comprised of lymphocytes, plasma cells and occasional heterophils, were also present. In experimentally infected ducks, lesions were similar with those observed in naturally infected birds.

**Figure 1.**
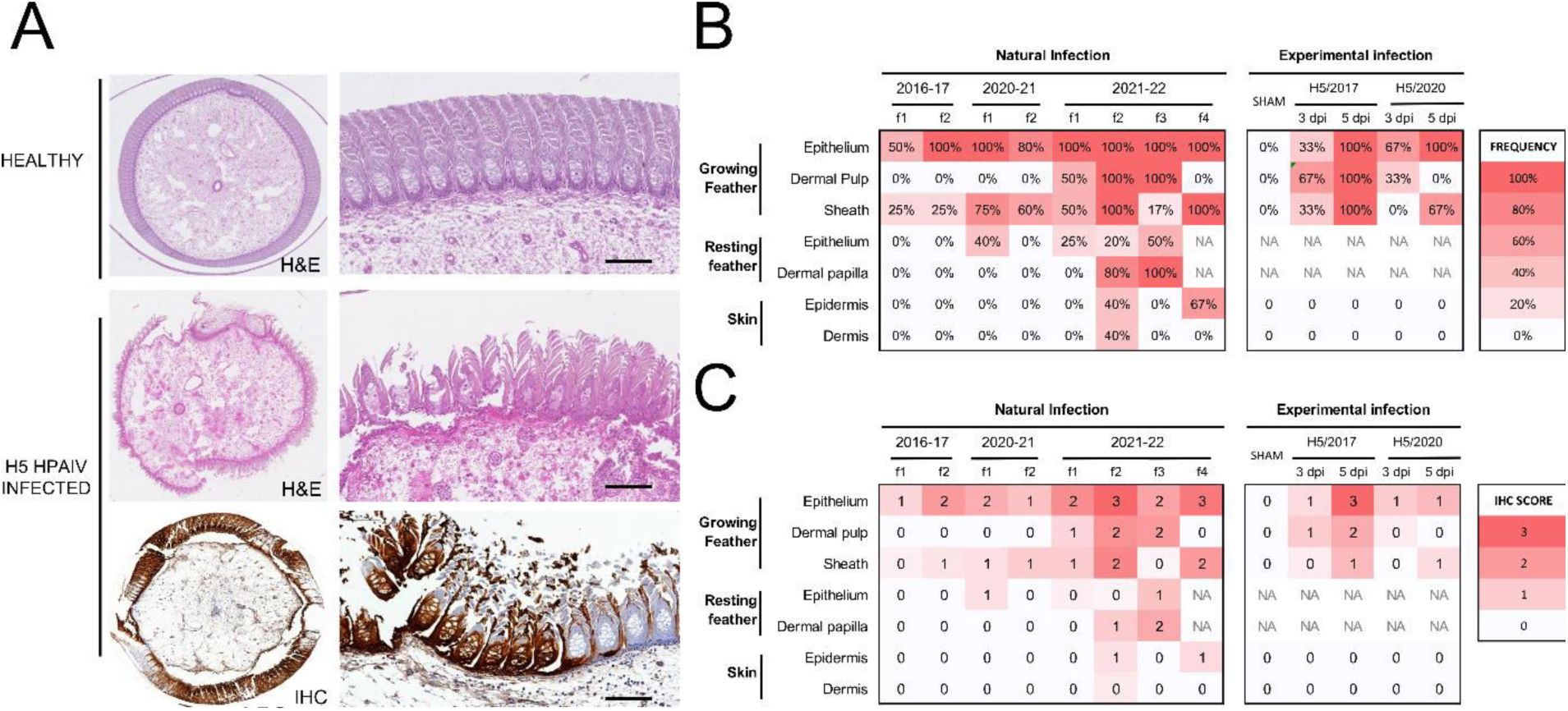
Feather lesions and *in situ* viral detection in commercial ducks naturally and experimentally infected with clade 2.3.4.4b H5 HPAIVs. (A), growing feather from a healthy duck showing normal dermal pulp surrounded by intact feather epithelium (H&E, top figure). In contrast, severe epithelial necrosis and disruption associated with inflammation of feather pulp (H&E, middle figure) and viral antigen detection (IHC, bottom figure) is observed in a growing feather from a duck naturally and experimentally infected with H5 HPAIVs, respectively. Bars represent 100 µm. Frequency (B) and average score (C) of viral antigen detection by IHC according to tissue compartment in naturally and experimentally infected ducks. Flocks are labelled as f and are numbered for each period. Flocks included 4-6 birds per flock.

### H5 HPAIVs have marked and persistent feather epitheliotropism

Viral antigen was detected within lesions in both cytoplasm and nuclei of cells (**Figure 1A**). Overall, in domestic ducks the most frequently and severely affected tegumental compartment was the growing feather epithelium, followed by the feather sheath (**Figure 1B, 1C**). In growing feather epithelium, viral antigen was frequently and extensively detected in barbules cells, barb cells (ramus), rachidial cells, and supportive cells from marginal and axial plates. In contrast, viral antigen detection was less frequent and widely spread in feather dermal pulp and resting feathers, except in ducks naturally infected with H5N1 from 2021-22. In the dermal pulp, viral NP was mainly detected in mesenchymal stromal cells and leukocytes. In resting feathers, viral antigen was inconstantly detected in dermal papilla and lining epithelium. Feathered skin dermis and epidermis were mostly negative, except in some ducks naturally infected with H5N1 from 2021-22. Viral antigen detection in other species including chicken, turkey, quail, geese, and wild swan is presented in **supplementary data 3**.

These results indicate that H5 HPAIVs have a marked tropism for feather epithelium in domestic ducks.

### H5 HPAIVs are detected in growing feathers early in the course of infection

In experimentally infected ducks, viral RNA was detected in growing feathers as early as 2 dpi, with peak of detection at 4 dpi and positive detection up to 14 dpi in the remaining ducks (**Figure 2A**). At 3 dpi, when the majority of the ducks were in the pre-symptomatic phase, viral antigens were multifocally detected within the dermal pulp and basal layer of growing epithelium, extending through marginal plates, and then in the lower portion of barb and barbule cells (**Figure 2B**). At 5 dpi, when mortality peaked, the detection of severe and diffuse viral antigens extended to barbs, barbules, and inter-barbular spaces (**Figure 2B**). Viral antigen distribution was similar in H5N8/2017 and H5N8/2020 infected ducks within feather tissue. However, immunoreactive areas tended to be more widespread in H5N8/2017 infected ducks.

**Figure 2.**
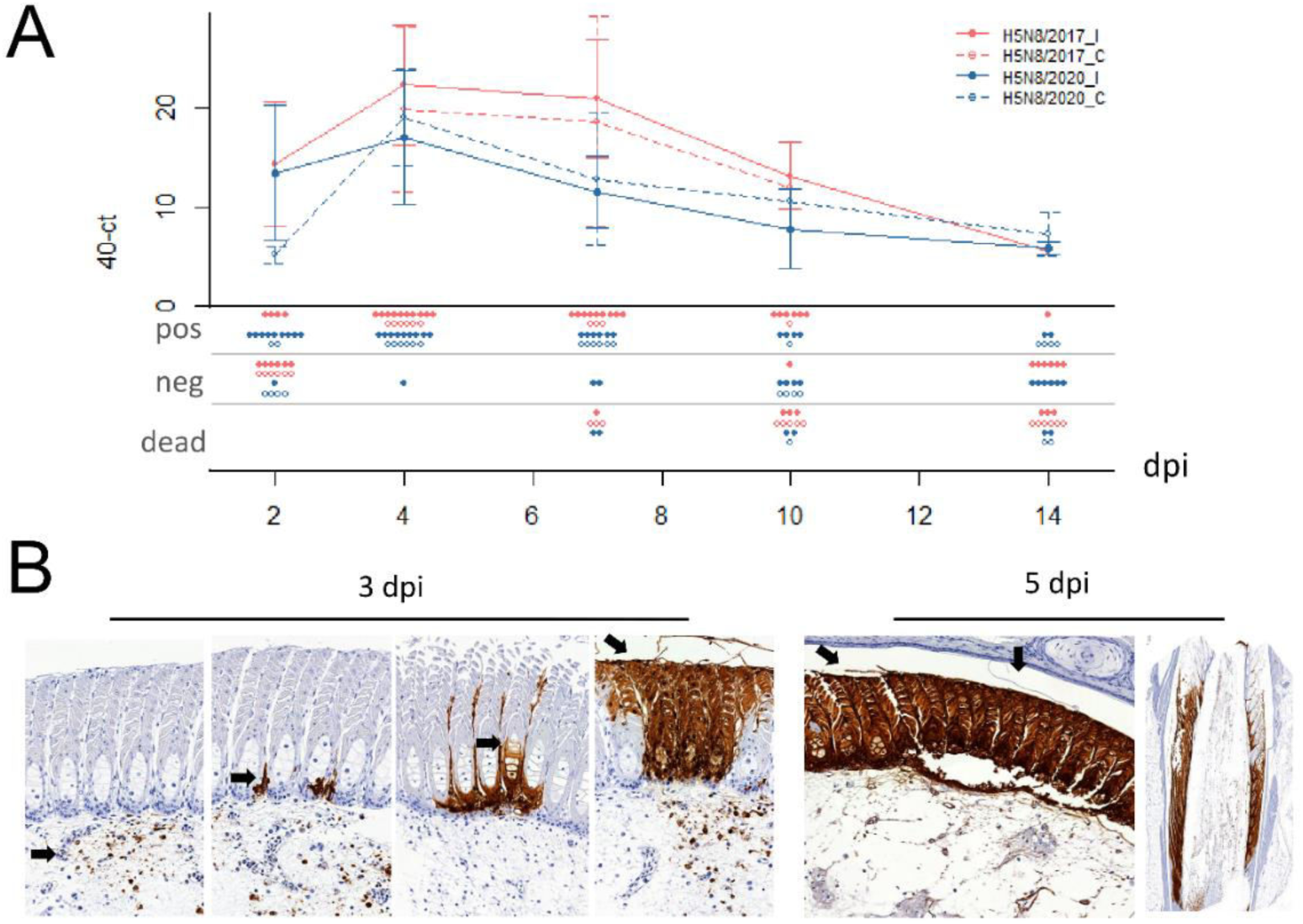
Viral antigen dynamics in commercial ducks experimentally infected with H5 HPAIV from 2017 and 2020. (A) Viral RNA detections in growing feathers from ducks experimentally infected with H5N8 HPAIVs from 2017 (red, H5N8/2017) and 2020 (blue, H5N8/2020), represented as 40-Ct values (>40 were considered negative). Means calculated on living birds at the time of sampling are represented as points, and standard deviations as hooks for both inoculated (I) and contact (C) birds. N+, N-, ND dots represent number of positive, negative and dead birds, respectively. H5N8/2017_I: mule ducks inoculated via the intrachoanal route with 5 log_10_ EID_50_ of H5N8/2017, H5N8/2017_C: mule ducks placed in contact with H5N8/2017 inoculated ducks 1 dpi, H5N8/2020_I: mule ducks inoculated via the intrachoanal route with 10^5^ EID_50_ of H5N8/2020, H5N8_C: mule ducks placed in contact with H5N8/2020 inoculated ducks 1 dpi. (B) Viral antigen detection in growing feathers of experimentally infected ducks. At 3 dpi viral antigens can be multifocally detected in the dermal pulp, extending to the marginal plate, followed by barb and barbule cells (arrows). At 5 dpi, viral antigens diffusely extend into the feather epithelium, anti-IAV NP IHC.

These results indicate that H5 HPAIVs rapidly infect and diffuse into the feather epithelium over the course of infection.

### Virion-shaped particles are observed in infected feather epithelium

Ultrastructural analysis was performed on a total of six different sections of growing feathers obtained from the caudal tract of a duck experimentally infected with H5N8/2017. Regions of interest were selected based on semi-thin sections and observation of epithelial necrosis. Filamentous particles, ranging 60-80 nm in diameter and up to 500 nm in length, were seen budding from the cytoplasmic membrane of the feather epithelium and accumulating between barbules, extracellularly (**Figure 3**).

**Figure 3.**
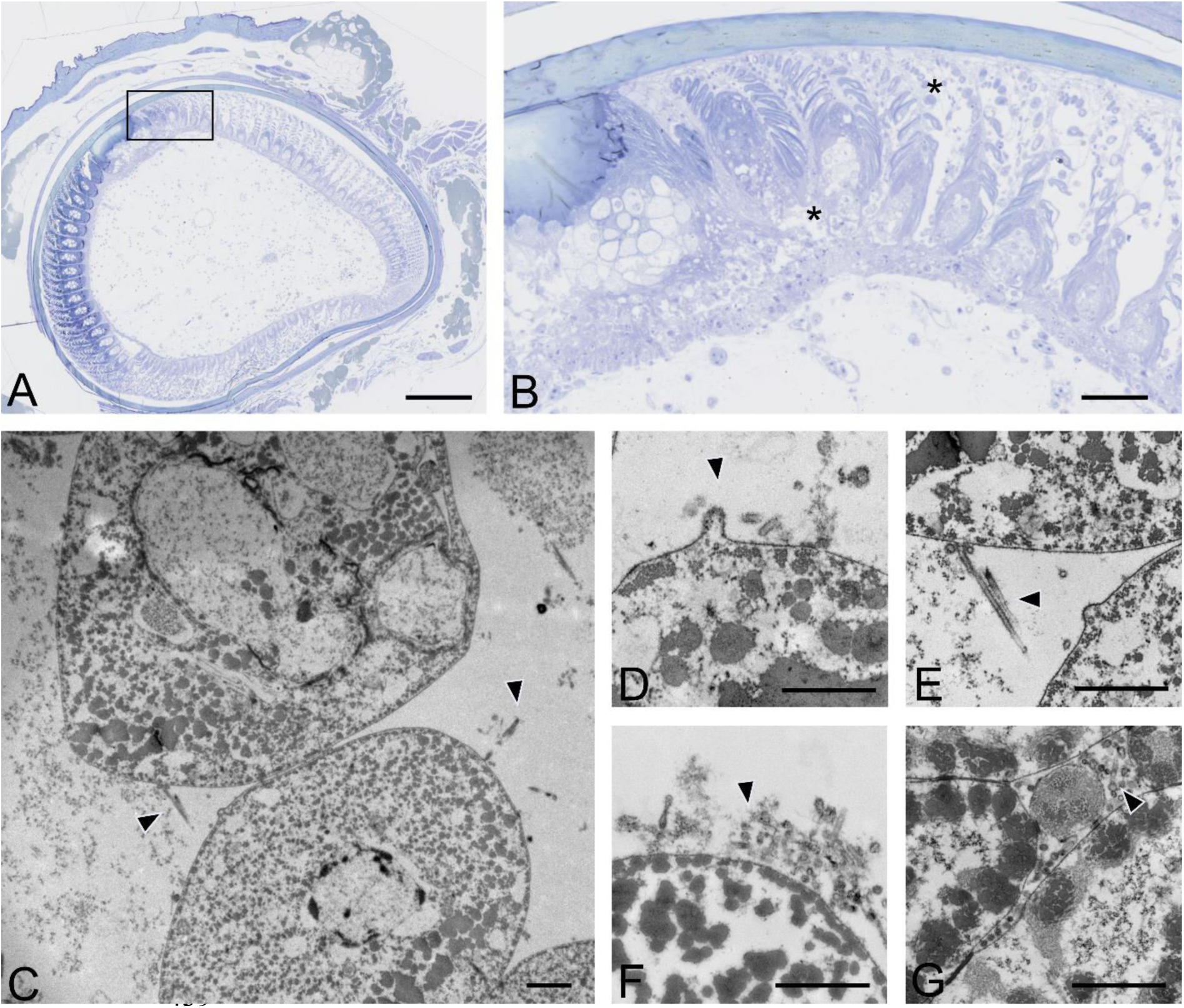
Ultrastructural viral detection in growing feather epithelium of commercial ducks experimentally infected with H5 HPAIVs. (A) Ultrastructural images of feather epithelium of a duck experimentally infected with H5N8/2017 HPAIV at 6 dpi was assessed after selecting regions of interest (ROI) on histological sections. Bar, 200µm. (B) ROI within feather growing epithelium included barbules, inter-barbular spaces, and necrotic areas. Bar, 50µm. C-G, Detection of budding and free, filamentous virion-shaped particles (arrowheads) within intercellular spaces of differentiating barbules. Transmission electron microscopy. Bar, 500 nm.

These results indicate that infection of the feather epithelium generates production of virions released extracellularly between barbules.

### High titers of infectious material are associated with immature feather epithelium

Both the detection of viral RNA and proteins in the feather epithelium and the observation of particles evocative of filamentous influenza virions, prompted us to determine whether the infected feather epithelium produced infectious particles. To do so, immature duck feathers from naturally infected flocks were carefully dissected after sheath disinfection (**Figure 4A**). The association of infectious material with the various feather subcompartments was then determined by viral titration in MDCK cells (**Figure 4B**). Although moderate levels (mean concentration of 2.16 log_10_ FFU/mL) of infectious material were present on feather sheaths, ethanol incubation led to a drastic reduction of infectivity, below the assay detection limit, in 8/10 samples, ruling out subsequent contamination of internal feather elements by external infectious material (**Figure 4B**). High concentrations of infectious material (mean concentration around 4.39 log_10_ FFU/mL) were detected after simple incubation, without agitation of epithelium samples. Similar concentrations (mean concentration of 4.39 log_10_ FFU/mL) were detected after replacement of the diluent and vortexing. Finally, mean concentrations of 3.53 log_10_ FFU/mL were detected in feather pulp sampled in the area closest to the skin. Overall, viral titers were statistically higher in the epithelial compartment compared to the pulp and sheath compartments (p<0.05).

**Figure 4.**
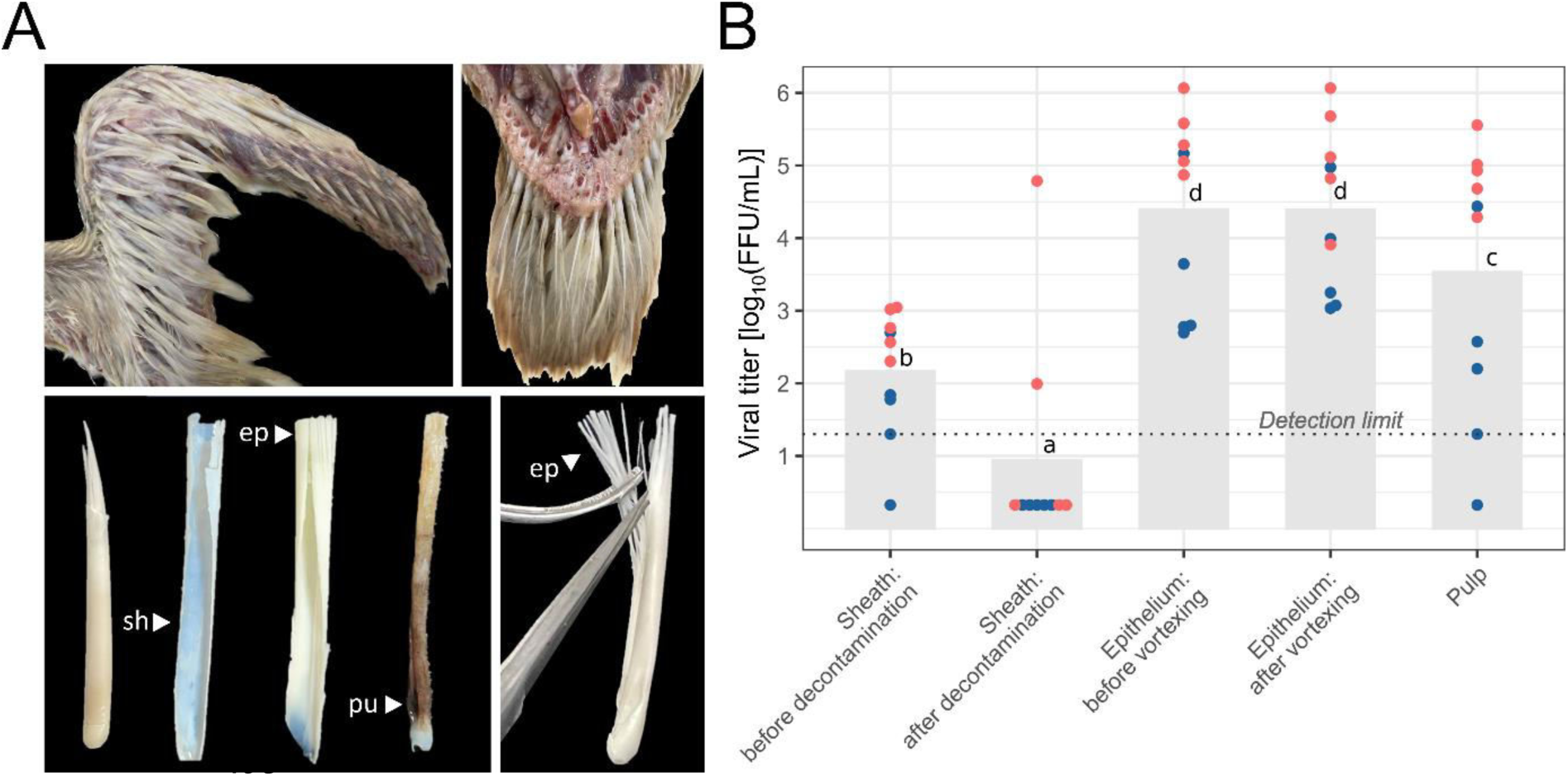
Viral infectivity of growing feathers in commercial ducks naturally infected with clade 2.3.4.4b H5N1 HPAIV. (A) Numerous growing feathers can be observed in the wing (top left photo) and caudal tract (top right photo) of a 40-day-old male mule duck at necropsy. Detached growing feathers are comprised of an external outer sheath (sh) and a growing and differentiating epithelium (ep) supported by a central core of dermal pulp (pu) (bottom left photo). For viral titration, distal epithelium was collected (bottom right photo). (B) Viral titers in MDCK cells, expressed as log_10_ FFU/mL, determined on feather outer sheath before and after decontamination, feather epithelium with and without mechanical disruption, and feather pulp. Histograms, dotted line, colors and dots represent viral mean titers, limit of detection, the two flocks included in the analysis, and subjects, respectively. Different letters indicate significant difference at p<0.05.

These results support the hypothesis that infectious viral particles are produced by the infected immature feather epithelium.

### Feather fragments and viral antigens colocalize in dust from influenza outbreak farms

Histologically, dust collected from four H5 HPAIV-positive farms revealed the presence of frequent light to dense, eosinophilic and elongated structures admixed with bacteria, fungi and granular material of unknown origin (**Figure 5**). Prior to IF, Corneous beta-protein (CBP) antigen detection was first determined by IHC on normal feathered skin: CBP was detected in cornifying cells, including barbules and the feather outer sheath (**supplementary data 4**). Subsequently, double IF, targeting both viral IAV NP and CBP, was conducted on sections of feathered skin of a duck experimentally infected with H5N8/2017 to assess positive detection of both targets, and then on dust blocks obtained from H5 HPAIV-positive farms. In feathered skin, both CBP and IAV NP antigens colocalized in the growing feather epithelium, particularly in the barbule cells (**Figure 6**). IAV NP antigen was also observed between barbules. In dust blocks, CBP antigen was detected within the elongated structures. AIV NP antigen colocalized with the CBP-positive elongated structures and the granular material (**Figure 6, supplementary data 5 and 6**).

**Figure 5.**
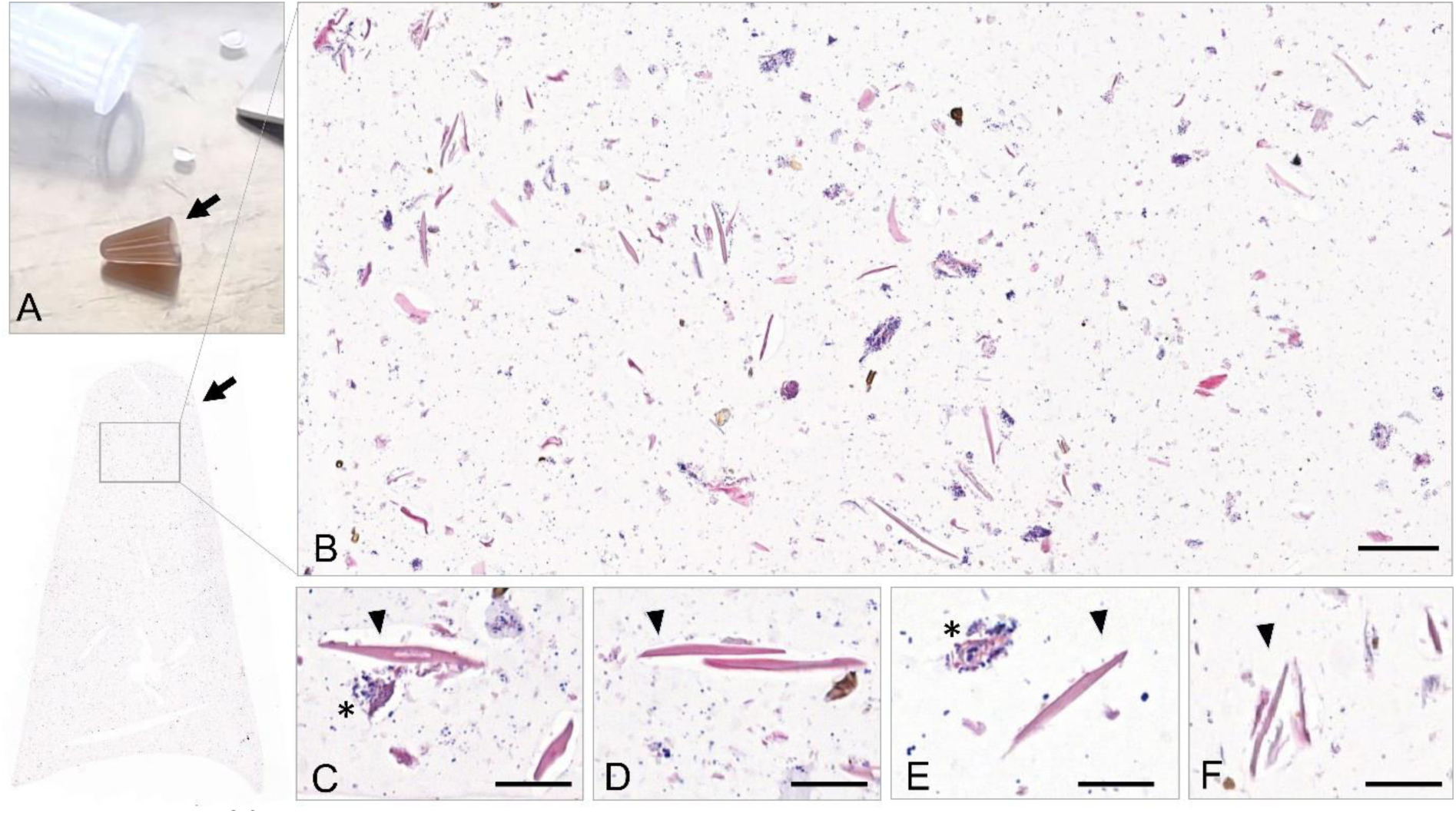
Histological analysis of aerosols (dust) collected from H5 HPAIV-positive farms. (A) Histogel-based Coriolis Compact (Bertin Technologies) sampled dust blocks (arrows). (B-F) Dust blocks from H5 HPAIV-positive farms stained with H&E revealed the presence of frequent eosinophilic elongated structures (arrowheads), admixed with bacteria (asterisks), fungi and granular material of unknown origin. Bar, 50µm (A) and 20µm (B).

**Figure 6.**
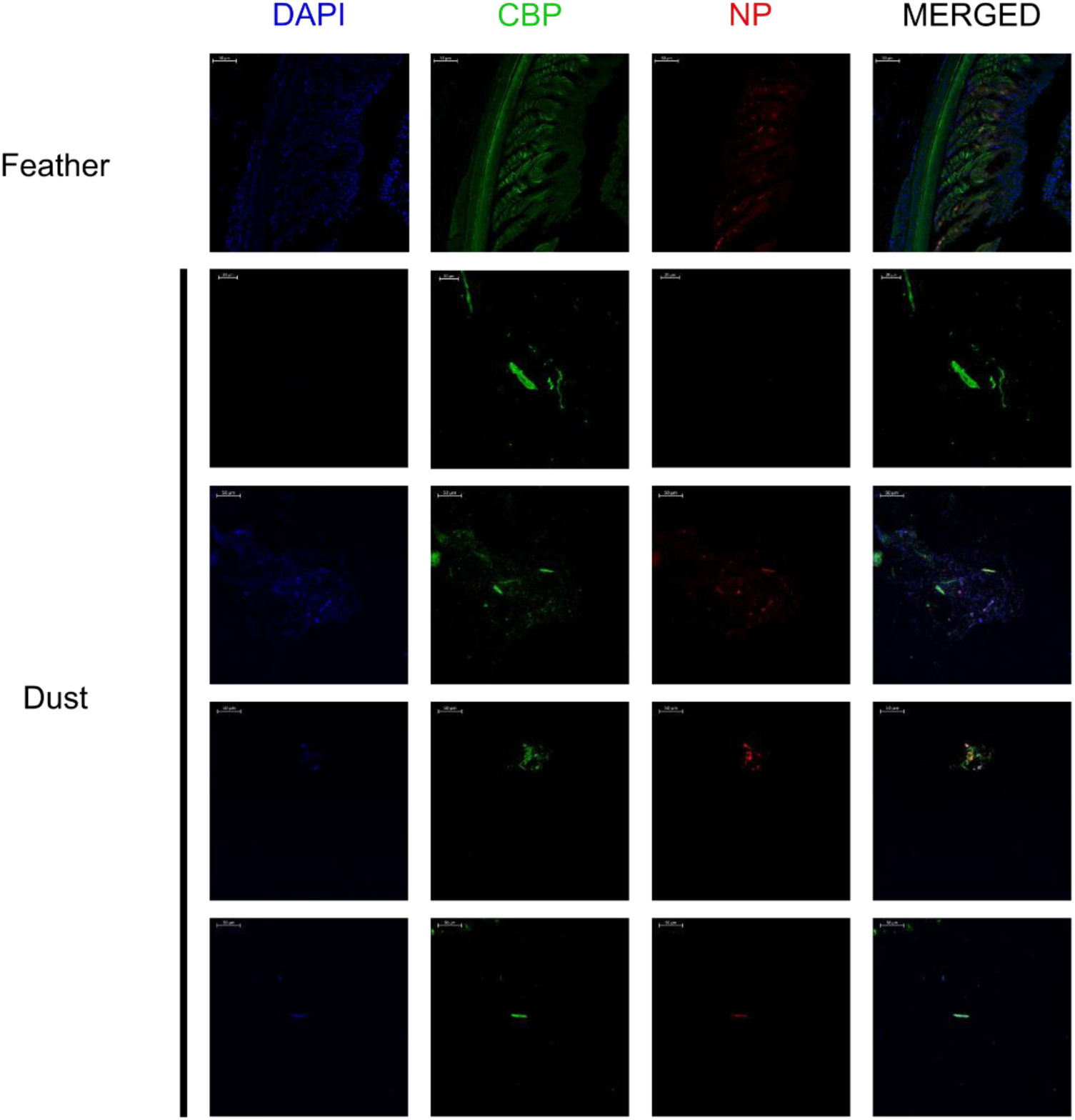
Immunofluorescent detection of avian CBP and influenza A NP antigen in growing feather and aerosols (dust). Immunofluorescent detection of avian CBP and influenza A NP antigen in a growing feather of an H5N8/2017 experimentally infected mule duck, and in histogel-based dust block sampled by Coriolis Compact (Bertin Technologies) from four HPAIV-positive farms during H5N1/2021 French outbreaks. 4’,6-diamino-2-phenilidole (DAPI, blue), Corneous beta-protein (CBP, green), Nucleoprotein influenza A (NP, red).

These results indicate that fragments detached from the infected feather epithelium are aerosolized in infected poultry houses and remain infected.

## Discussion

The extreme contagiousness of avian influenza viruses and, in particular, of HPAIVs derived from the Gs/GD lineage, is now universally recognized, especially in ducks. This is due to both their high susceptibility to viral infection and their great shedding capacity. Experimental studies showed that both oropharyngeal and cloacal viral shedding occur as early as a few hours post-infection and can last up to 14 days^5, 31–33^. In comparison, the duration of viral excretion in infected chickens is much shorter (<2 days) due to their short MDTs, contributing to the higher excretion potential of ducks compared to gallinaceous birds^4, 5, 24^.

Feather tropism of HPAIVs is now well documented, particularly for viruses of the Gs/GD lineage^7–9, 32, 34^. However, virological investigation of feather tropism have mainly focused on the feather pulp, regarded both as a marker of systemic infection and a target for diagnostic purposes, but not on the feather epithelium, as a potential excretion route of infectious viruses. In these studies, detached feather follicles were homogenized following extraction and processed as a whole biological matrix. As a consequence, the precise tissue location was lost^8, 12, 32, 35^. In our study, particular attention was given to the dissection of growing feathers, allowing to differentiate viral titers obtained from the feather epithelium, pulp, and sheath (**Figure 4**).

Our data show a marked tropism of clade 2.3.4.4b H5 viruses for the epithelial structures of growing feathers in commercial ducks. A similar marked epitheliotropism was identified in other waterfowl species, including commercial geese and wild mute swans naturally infected with H5 HPAIVs (**supplementary data 3**). On the contrary, in gallinaceous birds naturally infected with H5 HPAIVs, including commercial chickens, quail, and turkeys, viral antigen detection was more frequent and widely spread in the feather pulp and skin dermis compared to the feather epithelium (**supplementary data 3**). These results are consistent with experimental infections^9^ and suggest rapid death of infected galliformes is associated with limited viral extension from pulp to the feather epithelium.

The total number of feathers present in a single bird is poorly known although, on average, it could account for 3-6% of the total body weight of an adult individual^36^. Attempts to manually count feathers in individual birds have been reported in the 20^th^ century. In 1937, George Andrew Ammann allegedly counted 25,216 feathers in a swan. Phoebe Knappen reported a total of 11,903 feathers in a female adult duck^37^. In young birds, feather growth is an extremely fast and relatively synchronous process, that can reach up to 4 cm over 10 days in 40-day-old ducks^38^. For this reason, the feather epithelium compartment represents an extremely large surface area and volume in young individuals (**Figure 4A**). In contrast, adult birds have a higher number of mature feathers that are associated with lower viral detection.^32^ However, molting can result in increased proportion of growing feathers in adults. Different periodic and spontaneous molting processes can be observed in wild birds, which can be influenced by photoperiod, nutritional deficiency, migration timing constraints, and global warming^39, 40^. In the poultry industry, molting can be induced to increase egg production and quality in laying chickens^41^. Therefore, attention should not only be given to the study of host species but also to host age, breeding practices, plumage molt, and their potential impact on viral dissemination.

Normal and pathological behaviors may also lead to increased viral transmission from birds with infected feathers to other individuals. Ducks are well known for their preening behavior, which is very specific of waterfowl and to which they devote several hours a day. This behavior can involve only one bird (self-preening) or several congeners (allo-preening) ^33^. Additionally, feather pecking is a pathological injurious behavior resulting in removal and eating of feather’s congeners. Feather picking is a self-inflected chowing, biting, and plucking of feathers that can be seen when birds are reared in confined and poor environments^42^. Consequently, feathers represent an external organ strongly involved in normal and pathological ethograms of wild and domestic birds. This suggest possible direct transmission and ingestion of viral particles from feathers.

The quantitative importance of the feather excretion route, compared to the respiratory and digestive shedding routes, still needs to be assessed^33^. Nevertheless, although a definitive conclusion is still premature, the intensity of the viral signal detected in the feather fraction identified in dust samples is remarkable. Even though current data on the composition of poultry dust refer only to chickens and suggest that feather particles make up as much as 10% of the total mass of the dust present in poultry houses,^17^ a proxy to other species can be made, underlying the quantitative importance of animal exposure to this type of substrate. A clear parallel can be drawn between AIVs and the etiological agent of Marek’s disease in chickens; Gallid alphaherpesvirus 2 replicates to high titers in the epithelium of the feather follicles, and mature virions are released in association with debris originating from the epithelial layer. In this form, herpesvirus can survive for several months in the poultry house, which is much longer than expected for a herpesvirus^43^. A similar excretion route was found for the chicken infectious anemia virus^44^.

Staining experiments on dust samples collected from the environment of infected duck farms confirmed that viral antigens are mostly co-detected with avian CBPs. In feathered skin, CBP antigens could be detected in barbs, barbules, feather sheath, and the outermost layer of epidermis, but viral antigen detection was mostly positive in feather epithelium. Assessing the infectivity of organic material released into the environment is always a challenge. In order to overcome this issue, we tested the infectivity of the epithelial layer in a controlled setting. This experiment showed that the infectious viral load was higher in the epithelial layer compared to the pulp and feather sheath, supported by the observation of filamentous virion-shaped particles by electron microscopy in infected feather epithelium. Filamentous influenza viral particles are indeed preferentially observed after infection of epithelial cells and their genesis appears to depend on both host and virus-derived factors.^45^ The relative frequency of filamentous versus spherical viral particles produced by infected feather epithelium is currently under investigation. Altogether, these data support the notion that infected duck feather debris are infectious. The persistence of viral infectivity in the environment remains to be assessed, in particular for long-distance contamination and between-farm dissemination.

Active viral shedding occurs along the time course of infection from pre-clinical to clinical phases^4^. However, attention should also be given to the passive post-mortem viral shedding from infected carcasses littering the ground. Stability in detached infected feathers has been demonstrated for both H5 and H7 HPAIVs in experimental conditions^12, 15^. Feathers can remain infectious for up to 15 days at 20°C and 160 days at 4°C for H5N1 HPAIV^15^. In addition, during the decomposition process, epidermal slippage and hair loss can occur in mammals during the early decay stage (24h and 24 days post-mortem) ^46^. In birds, feather loss and disruption take place throughout the decomposition process,^47^ underlying the importance of rapid disposal of infected carcasses and complete environmental disinfection, considering also the risk of exposing cadaveric fauna (insects, arthropods etc), scavenger birds, and mammals to infectious tegument. Dispersion of feather debris may also occur during transportation of ducks from the farm to the slaughterhouse, during processing, or in live poultry markets, which play a critical role in the epidemiology and evolution dynamics of HPAIVs in many low-and middle-income countries^48^.

By identifying another route of viral shedding for ducks, these results open the way to a paradigm shift in the epidemiology of avian influenza. Transmissibility is influenced by viral shedding and environmental stability^6^. Virions may be protected from physical factors or even disinfectants, resulting in an unexpected persistence of infectivity in the environment. However, whether any particular resistance properties are associated with this route of excretion, thanks to the protection provided by cornified cells, endogenous or uropygial lipids, still needs to be clarified^16, 49, 50^.

Our findings support the importance of an underestimated transmission route in ducks, i.e., the infected plumage, in the environmental shedding and dissemination of H5 HPAIVs. Further investigations are needed to establish the importance of this route compared to the fecal-oral and the respiratory-aerosol routes, and to define its impact in the implementation of biosecurity measures.

## Acknowledgements

This study was performed in the framework of the “Chaire de Biosécurité et Santé Aviaires”, hosted by the National Veterinary College of Toulouse (ENVT) and funded by the Direction Generale de l’Alimentation, Ministère de l’Agriculture et de la Souveraineté Alimentaire, France. The animal experiment was partially funded by the Veterinary Biocontained facility Network (VetBioNet) [EU Grant Agreement INFRA-2016-1 N°731014]. The authors would like to thank Manel Vinyes (GALLSA) for providing the ducks for the experimental infections, and Miquel Nofrarías, Maria José Valdez, and Rosa Valle for their technical assistance. We thank Thomas Figueroa, Pierre Bessière and Romain Volmer for providing viral stock of reverse genetics-engineered virus. We would like to express our sincere gratitude to Professor Lorenzo Alibardi from the University of Bologna for providing primary antibody aliquotes for the detection of corneous beta proteins. We gratefully acknowledge Cell Imaging Facility of INFINITy for access and assistance to the confocal microscope. The authors are thankful to Ms Aurore Mathon and Ms Grace Delobel for the design of the graphical Abstract and the English proofreading, respectively.

## Author contributions

**N.G., F.F., J-L.G:** conception of the study**. N.G., F.F., K.B., S.S., N.M., J-L.G.:** conceptualization, discussion and writing. **NG**., **M.C**., **M.D**., **A.S., S.S., F.F., J-L.G**: pathological and virological investigations. **N.G., F.F., K.B., A.P., G.C., N.M., J-L.G** : animal experiment**. B.P**: electron microscopy.

## Data availability statement

The authors confirm that the datasets generated during and/or analysed during the current study are available from the corresponding author on reasonable request.

---

**Supplementary data 1.**
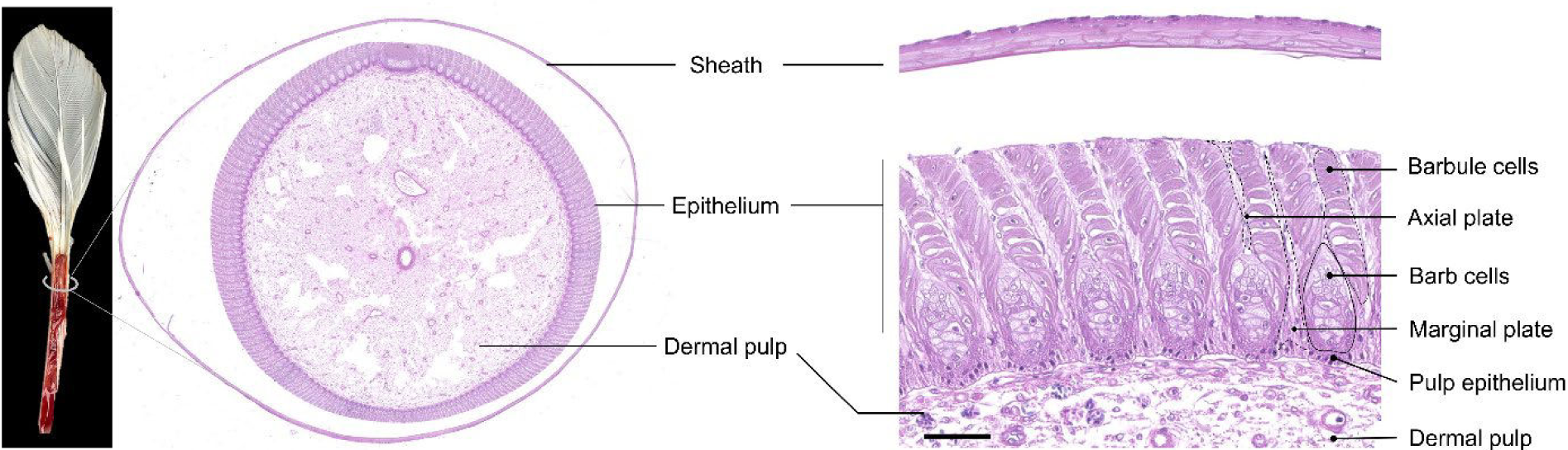
Histology of a growing feather. Growing feathers are surrounded by an outer sheath. The dermal pulp consists in a loose stromal tissue and blood vessels. The feather epithelium is composed of a basal pulp epithelium, differentiating (cornifying) barb cells (ramus) supporting numerous barbule cells, and supportive cells (axial and marginal plates) that will disintegrate by programmed cell-death. Additional information about functional histology and ultrastructure are provided by several references ^50–52^

**Supplementary data 2.**
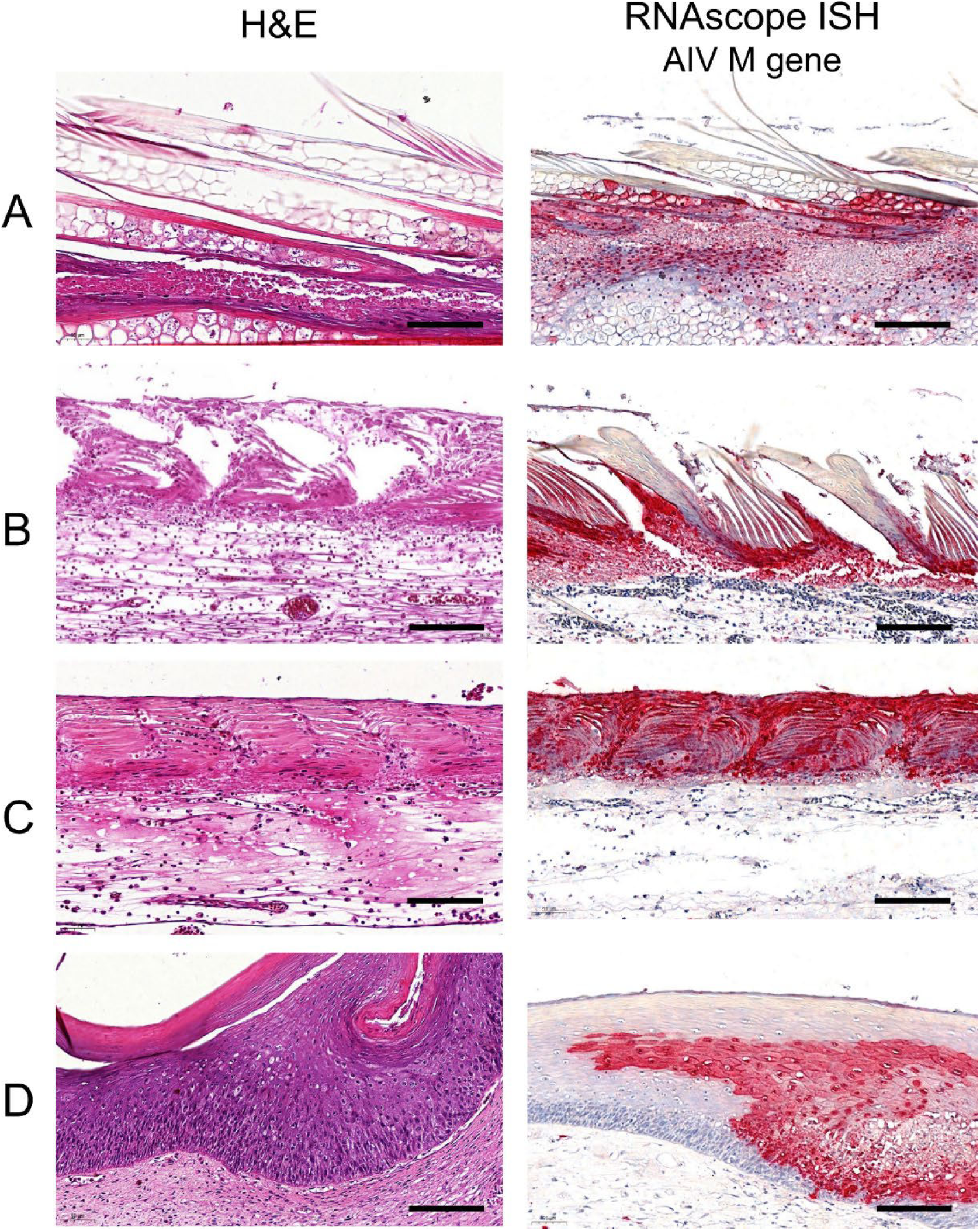
Proximo-distal lesions and viral RNA distribution in growing feathers of domestic ducks naturally infected with clade 2.3.4.4b H5 HPAIVs. Severe epithelial necrosis and disruption associated with widespread epithelial viral RNA detection in distal barbs and barbules (A); late barb ridges (B); early barb ridges (C); proximal epidermal collar (D). Hematoxylin and Eosin (H&E). RNAscope *in situ hybridization* targeting avian influenza virus *Matrix* (*M*) gene RNA (RNAscope ISH).

**Supplementary data 3.**
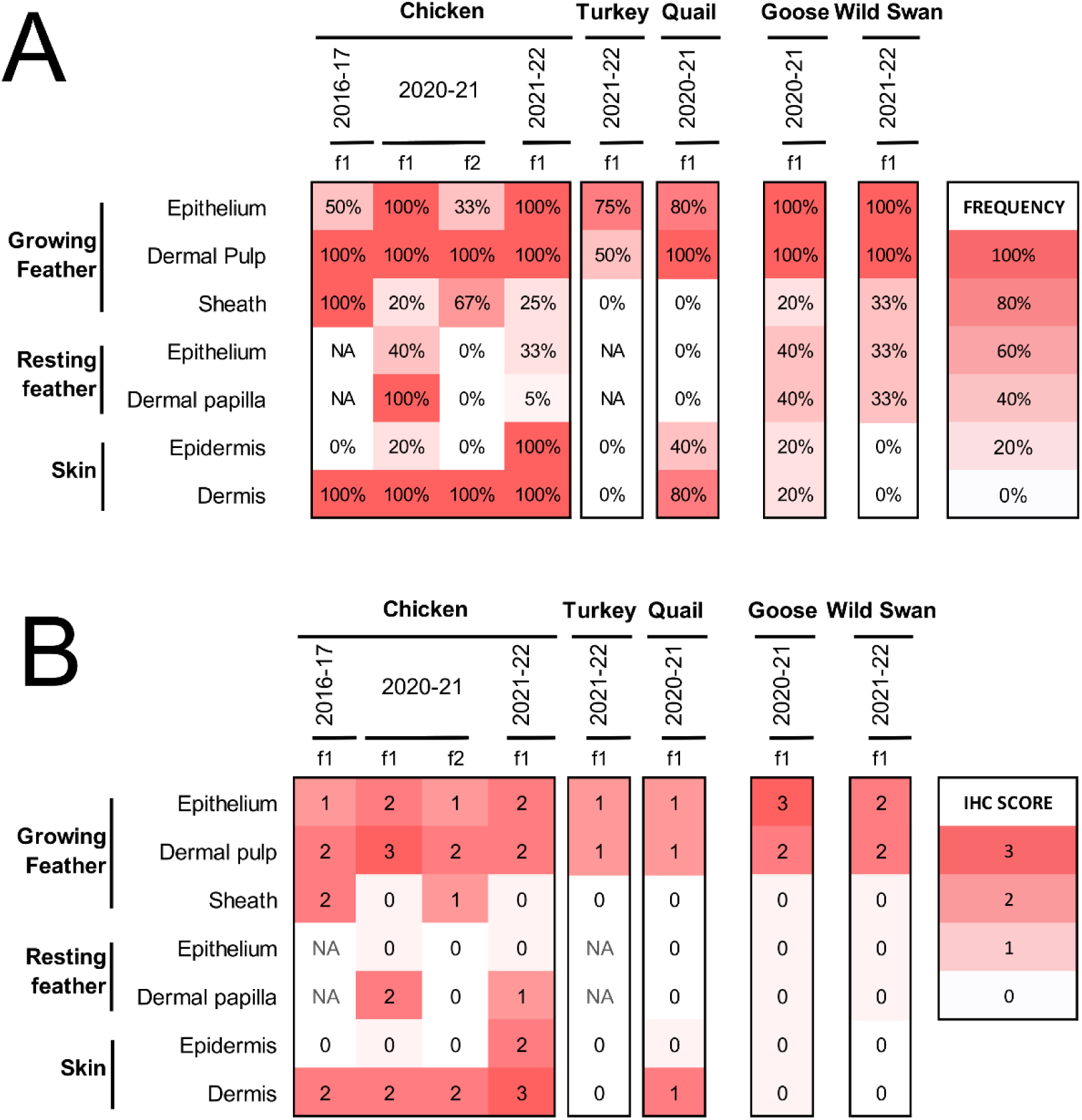
Viral *in situ* detection in commercial galliformes, geese, and wild waterfowl naturally infected with clade 2.3.4.4b H5 HPAIVs. Feathered skin was collected in naturally infected commercial chickens (4 flocks, n=3-5 birds per flock), turkeys (1 flock, n=4), quail (1 flock, n=5), geese (1 flock, n=6), and wild mute swans (n=3) found dead between 2016 and 2022. Histopathology as well as anti-nucleoprotein influenza A immunohistochemistry were performed similarly to duck samples (see material and methods). Table A and B represent frequency and average score of viral antigen detection, respectively, and according to tissue compartment. Flocks are represented as f and are numbered for each period.

**Supplementary data 4.**
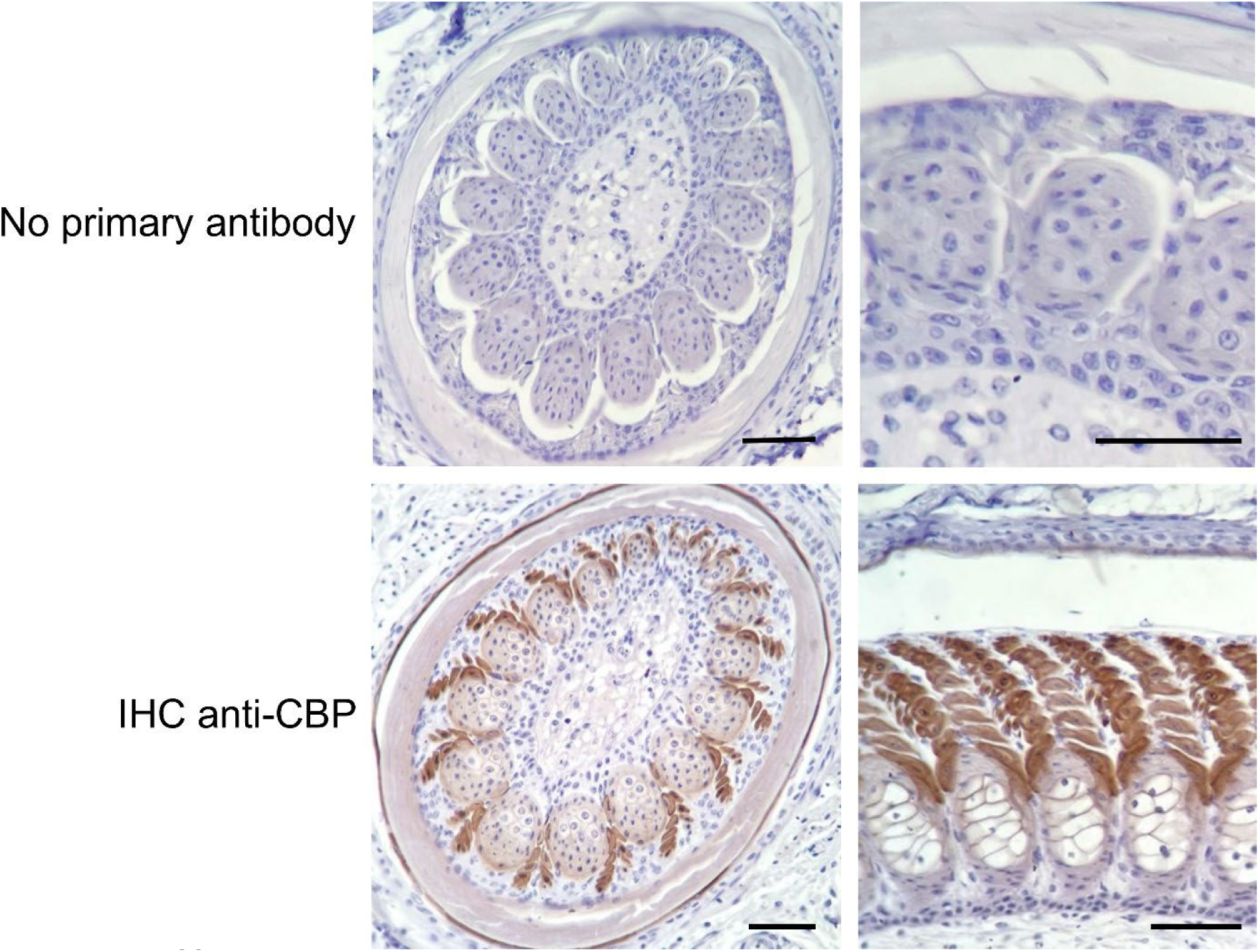
Immunohistochemistry targeting Corneous Beta-Proteins (CBP) in feathered skin. Top images represent negative control (no incubation of anti-CBP primary antibodies). Bottom images represent positive cytoplasmic antigen detection, marked in barbules cells, mild in barb cells. Antigen detection is also positive in the feather outer sheath. Bar, 50µm.

**Supplementary data 5.**
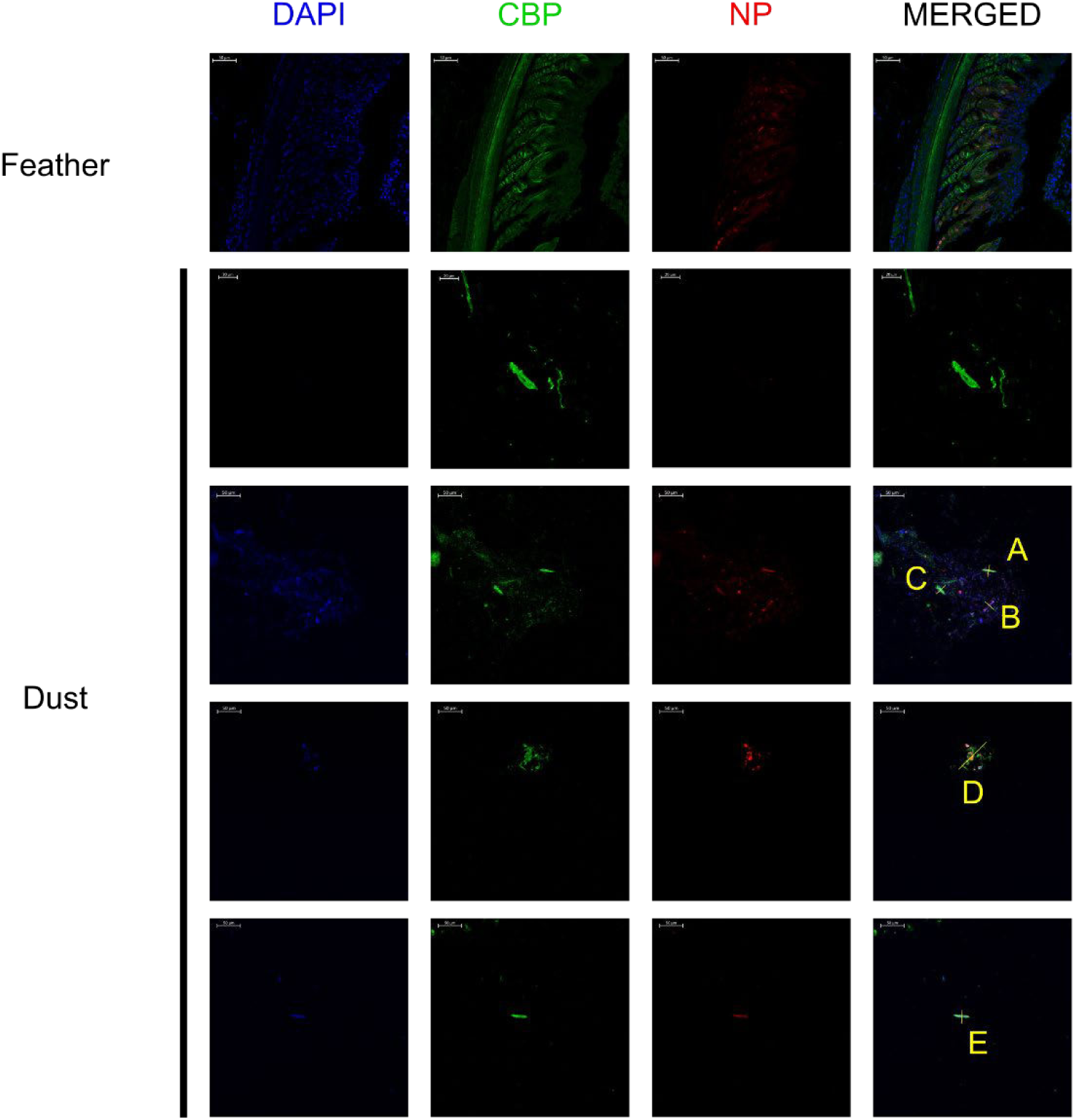
Immunofluorescent detection of avian CBP and influenza A NP antigen in both H5N8/2017 mule duck growing feather and histogel-based dust block sampled by Coriolis Compact (Bertin Technologies, https://www.bertin-instruments.com) in H5N1/2021 French outbreaks. Dapi (blue), Corneous beta-proteins (CBP, green), Nucleoprotein Influenza A (NP, red). Letters A-E indicate regions were colocalization graphs were assessed.

**Supplementary data 6.**
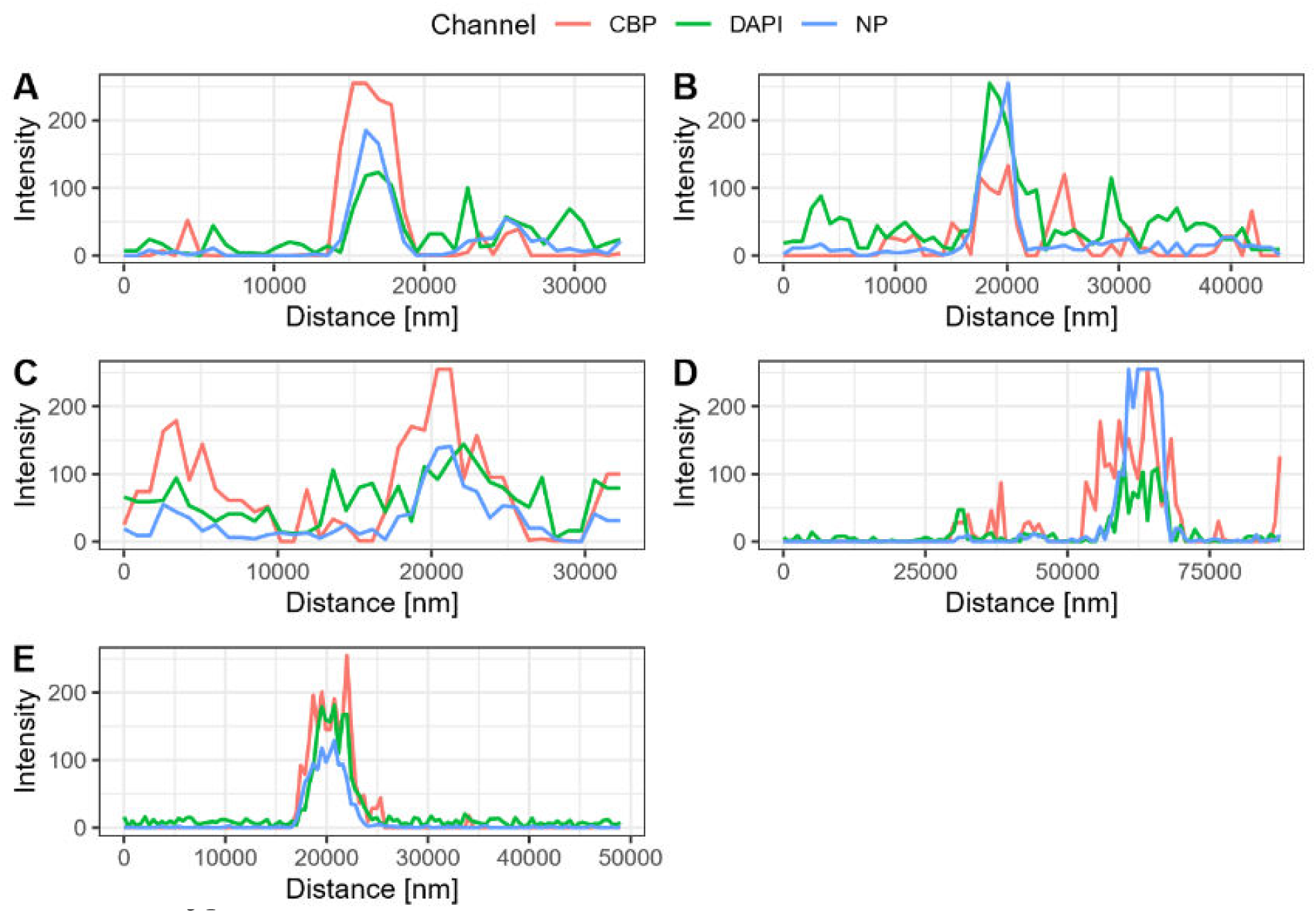
Immunofluorescent detection of avian CBP and Influenza A NP antigens in dust obtained from HPAIV-positive farms and growing feathers. Colocalization graphs of A, B, C, D, and E regions.

